# Rapid sperm capture: High-throughput flagellar waveform analysis

**DOI:** 10.1101/551267

**Authors:** M.T. Gallagher, G. Cupples, E.H. Ooi, J.C. Kirkman-Brown, D.J. Smith

## Abstract

Flagella are critical across all eukaryotic life, and the human sperm flagellum is crucial to natural fertility. Existing automated sperm diagnostics (CASA) rely on tracking the sperm head and extrapolating measures. We describe fully-automated tracking and analysis of flagellar movement for large cell numbers. The analysis is demonstrated on freely-motile cells in low and high viscosity fluids, and validated on published data of tethered cells undergoing pharmacological hyperactivation. Direct analysis of the flagellar beat reveals that the CASA measure ‘beat cross frequency’, does not measure beat frequency. A new measurement, track centroid speed, is validated as an accurate differentiator of progressive motility. Coupled with fluid mechanics codes, waveform data enables extraction of experimentally intractable quantities such as energy dissipation, disturbance of the surrounding medium and viscous stresses. We provide a powerful and accessible research tool, enabling connection of the cell’s mechanical activity to its motility and effect on its environment.

## Introduction

About 100 million men worldwide are suggested to be subfertile (Inhorn and Patrizio, 2015), but for many of them an accurate diagnostic cause remains elusive. The ability of sperm to successfully migrate through the female reproductive tract is key to natural fertilisation, however current techniques for assessing sperm motility lack diagnostic power and mechanistic insight. The heterogeneity of human sperm requires tools that can acquire and analyse (2018) and associated special issue). Such systems are widely used for veterinary work and in domestic animal breeding, conservation, and toxicology (Amann and Waberski, 2014), but have not yet made the breakthrough into routine clinical usage. The reasons for this have been discussed elsewhere (Gallagher et al., 2018b).

Standard CASA systems produce a motility assessment from the track of the head movement, however this does not enable causative or mechanistic insight because it lacks detail on the movement of the flagellum. Knowledge of this beat could provide hitherto untapped information for the estimation of non-visible attributes such as the contribution of different metabolic path-ways, and modulation in response to the physical and biochemical environments (Ooi et al., 2014). The capability to capture flagellar movements and associated mechanistic insights has broader applicability in the life sciences including the role of cilia in embryonic development (Smith et al., 2019), swimming of multiflagellate microorganisms (Wan and Goldstein, 2018), the use of high-speed holographic microscopy to image the flagellar waveforms of malaria parasites (Wilson et al., 2013), and even the design of hybrid bio-robots for biomedical applications (Xu et al., 2017).

The use of computers for flagellar tracking was pioneered by Hiramoto and Baba (1978), enabling the semi-automated capture of sperm flagellum (see Inaba and Shiba (2018) for an overview of the BohBoh system (Bo-hBohSoft, Toykyo, Japan)). This software has been used for a number of studies of sperm hyperactivation (Kaneko et al., 2007; Ohmuro and Ishijima, 2006), for example chemotaxis/chemokinesis (Guerrero et al., 2010; Shiba et al., 2006) and the movement of other flagellated species such as *Leischmania* (Mukhopadhyay and Dey, 2016; Reddy et al., 2017). Our own group’s previous work has employed bespoke semi-automated algorithms (combined with significant manual input) to enable to analysis of around 30–50 cells in studies of the effect of fluid rheology (Smith et al., 2009) and pharmacological stimulus (Ooi et al., 2014). Further recent methodological developments include the software of Walker and Wheeler (2018) and the SpermQ system (Hansen et al., 2018). We would however still characterise these techniques as requiring operator intervention (in particular manual segmentation) for the analysis of each set of imaging data, at least for the human sperm flagellum.

There remains therefore a need to progress flagellar analysis methodology further for both basic research and clinical applications; semi-automated analysis can only reasonably enable tens of cells to be analysed in a reasonable timeframe. This issue is particularly relevant to human sperm, which exhibit considerable heterogeneity (between cells, within cells over time, between ejaculates and between donors). It is therefore crucial to develop the tools to enable high-throughput flagellar analysis and thus take statistical measurements of flagellum movement over representative sample sizes.

This paper describes a free-to-use software package for high-throughput extraction and analysis of swimming sperm and their associated flagellar beat, FAST (**F**lagellar **A**nalysis and **S**perm **T**racking). Using FAST we have analysed 176 experimental microscopy videos and have tracked the head and flagellum of 205 progressive cells in diluted semen, 119 progressive cells in a high-viscosity medium, and 42 stuck cells in a low viscosity medium. The average time to analyse each video was a few seconds per cell on an Apple Macbook Pro. The package will be shown to provide positions in time of tracked sperm head and flagellum, and associated measurement of the flagellar arc-wavelength, flagellar beat frequency, and power dissipation by the flagellum in addition to providing the existing CASA measures. FAST is not designed to analyse raw semen, it is specifically for precise analysis of flagellar kinematics, as that is the promising area for computer use (Gallagher et al., 2018b). Flagellar capture will always require that cells are at a dilution where their paths do not frequently cross. This manuscript reports detailed statistics on flagellar arc-wavelength and beat frequency, the metabolic requirements of motility, as well as the relationship of these measures to the standard CASA motility measures. We will discuss the calculated velocity profiles for some characteristic sperm calculated using a recently-published open source fluid mechanics code (‘NEAREST’ Gallagher et al. (2018a); Gallagher and Smith (2018); Smith (2018)). Additionally, we introduce a novel measure TCS (track centroid speed) for differentiating progressive and non-progressive/immotile cells.

## Materials and Methods

Sperm flagellar movement exhibits major variations depending on the surrounding fluid environment and activation state. Analysis has been performed on two data sets: (1) a new data set on free-swimming cells in two different viscosity media, and (2) re-analysis of a previously-published dataset showing the response of tethered cells to a hyperactivating pharmacological stimulus. Details of the two data sets follow:

1. **Analysis of free swimming sperm in diluted semen and high viscosity media** Semen samples were provided by three unscreened normozoospermic donors (recruited at Birmingham Women’s and Children’s NHS Foundation Trust after giving informed consent). Semen samples were obtained through masturbation following 2–3 days’ abstinence. Cells were prepared according to two procedures in either diluted semen or high viscosity media. To prepare the diluted semen (DSM) samples were counted according to WHO guidelines (WHO, 2010) and diluted to a concentration of 10 M/ml in Earle’s Balanced Salt Solution (sEBSS) without phenol red, and supplemented with 2.5 mM Na pyruvate and 19 mM Na lactate (06-2010-03-1B; Biological Industries, Kibbutz Beit HaEmek, Israel), and 0.3% wt vol-1 charcoal delipidated bovine serum albumin (Sigma; A7906). For high viscosity media (HVM) cells were suspended in Earle’s Balanced Salt Solution (sEBSS) without phenol red, and with the addition of pyruvate, lactate, and bovine serum albumin, with the addition of 1% methylcellulose (M0512, Sigma-Aldrich, Poole, UK, specified so that an aqueous 2% solution gives a nominal viscosity of 4000 centipoise or 4 Pa s at 20°C). For DSM cells were imaged in a 10 μm depth chamber, while HVM was loaded by capillary action into flat-sided borosilicate capillary tubes (VITROTUBES, 2540, Composite Metal Services, Ilkley, UK) with length 50 mm, and inner dimensions 4 mm ×0.4 mm. One end of the tube was sealed with CRISTASEAL (Hawksley, Sussex, UK, #01503-00). Cells were selected for imaging by immersing one end of the capillary tube into a 1.5 ml Beem capsule (Agar Scientific, UK) containing a 200 μl aliquot of raw semen, within 30 minutes of sample production. Incubation was performed for 30 minutes at 37°C in 6% CO2. Both sets of cells were imaged using a Nikon (Eclipse 80i) microscope and negative phase contrast microscopy (objectives 10× 0.25 Ph1 BM ∞/– WD 7.0; and 20× 0.40 Ph1 BM ∞/0.17 WD 1.2), using a Basler Microscopy ace camera (acA 1300-200uc) at 169.2 frames per second with pixel size 4.8 μm ×4.8 μm, streaming data directly to a Dell XPS laptop using Pylon Viewer (v.5.0.11.10913, Basler). For DSM, the cells were imaged in 10 μm analysis chambers (10-01-04-B-CE; Leja Products B.V., Nieuw-Vennep, The Netherlands) at 10× magnification. Cells in high viscosity media were imaged at 2 cm migration distance into the capillary tube, and in the surface accumulated layer 10–20 μm from the inner surface of the capillary tube at both 10× and 20× magnification.
2. **Re-analysis of adhered cells under pharmacological stimulus** For full details see Ooi et al. (2014); briefly, cells adhered to a surface coated with 0.1% poly-D-lysine in a bespoke chamber filled with supplemented saline solution were imaged at 20× magnification with negative phase contrast, before and after perfusion of 4AP to induce hyperactivation. Images were captured at 332 Hz using a Hamamatsu Photonics C9300CCD. For each of experimental sets (1) and (2) the resulting imaging data was run through the FAST software package for tracking and analysis. In addition to the usual CASA kinematics obtained by tracking the position of the moving sperm head, FAST provides measurement of the flagellar arc-wavespeed, arc-wavelength (λ) and beat frequency (*f*).

### Computing resources and optics

FAST has been designed to work on Microsoft Windows, Apple Mac OS X, and Linux-based operating systems, with minimal hardware requirements. All analysis for the present manuscript were performed on a machine with a quad-core Intel i7 processor and 16GB of RAM although this is not the minimum required. FAST is designed to detect sperm contained in either AVI video files or TIF image stacks, with any imaging modality where the entirety of the sperm cell is bright on a dark background.

### Flagellar kinematic parameter calculations

FAST characterises the flagellar wave in terms of the tangent angle θ (*s, t*), a function of both the arclength along the flagellum *s* (measured in micrometers from the proximal end to the distal) and time *t*. We can then calculate the flagellar curvature as κ (*s, t*) = ∂θ/∂*s*, from which the beat frequency (*f*) and arc-wavespeed (*c*) can be derived, where we restrict the analysis region to be the section of the flagellum with 10 μm < *s* < 30 μm as a region which ensures the wave has developed, is large enough to contain all the relevant information, and is small enough to allow for fair comparisons between all cells.

#### Flagellar beat frequency

The flagellar beat frequency (*f*) is calculated by tracking the number of turning points of curvature for a choice of *s*, dividing by the time period *T* and then halving. This calculation is repeated for a number of points in *s* to reduce the effects of noise, and the median is taken as the final value for *f*.

#### Flagellar arc-wavespeed

To calculate the flagellar arc-wavespeed we begin, as for the beat frequency, by calculating the times corresponding to maxima in the curvature in time at *s* = 10 μm. Iterating forward in *s* we follow the crest of each wave in time, achieving a set of (*s, t*) pairs for each tracked wave. To fit the wavespeed to these points in a robust manner we formulate a linear mixed effects model (lme), consisting of a wave (straight line in (*s, t*)) with fixed effects on the slope and additional random effects on the slope and intercept to account for sperm metabolic noise inducing differences between waves. The output from solving this lme model not only gives a result for arc-wavespeed and statistics about the goodness of fit, but also statistics about the random within-cell variation in wave speed.

#### Flagellar arc-wavelength

Having calculated the flagellar beat frequency (*f*) and arc-wavespeed (*c*) the flagellar arc-wavelength is then given by λ = *c/f*.

### Flagellar power dissipation calculations

Following Ooi et al. (2014), who applied the restive force theory of Gray and Hancock (1955) (later modified by Lighthill (1976)), we write the hydrodynamic force exerted by the sperm flagellum by unit length as

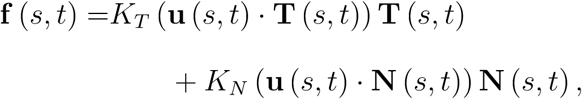

where **u** is the velocity of the flagellum at arclength *s* and time *t*, **T** and **N** are the unit tangent and normal vectors to the flagellum, and *K*_*T*_ and *K*_*N*_ are the tangential and normal resistance coefficients, where Lighthill’s ‘sub-optimal’ choice

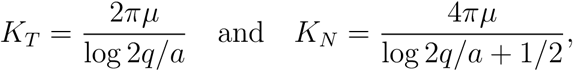

have been shown to be sufficiently accurate, where μ being fluid viscosity, *q.* = 0.09λ, and *a* being the typical radius of sperm flagellum. From this the power dissipation due to the beating of a section of flagellum with *s*_1_ *< s < s*_2_ can be calculated at a time *t* by

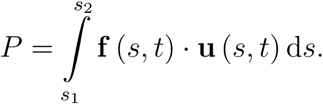

### CASA Measures

While the standard set of CASA measures have defined meanings, there are several differences in how the calculations are performed between many of the popular systems, not all of which are published. Here we will define each of the standard measures according to the WHO manual (WHO, 2010) and the methods by which FAST performs the calculations on head track locations **h**_*i*_ = (*x*_*i*_, *y*_*i*_) at times *t*_*i*_, at *n* + 1 points, *i* = 1,…,*n* + 1.

**VCL** time averaged curvilinear velocity in *μ*m/s:

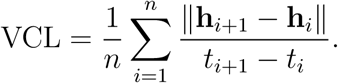

**VSL** straight-line velocity in *μ*m/s:

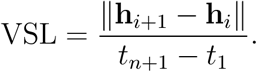

**VAP** average path velocity in *μ*m/s. In FAST we construct a robust average path by calculating a wavelet decomposition at level 7 of the path h using a discrete Meyer wavelet. We then discard the detail levels 1 - 4, and rebuild the path using wavelet reconstruction, giving the points A_*i*_.

**ALH_max_** maximum amplitude of lateral head displacement in *μ*m. Maximum displacement of the head track about the average path:

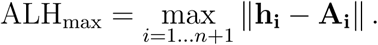

**ALH_avg_** average amplitude of lateral head displacement in *μ*m:

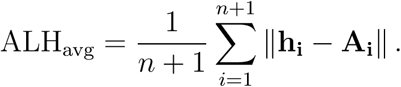

**LIN** linearity of the curvilinear path. LIN = VSL / VCL.

**WOB** wobble, a measure of oscillation of the curvilinear path about the average path. WOB = VAP / VCL.

**STR** straightness of the average path. STR = VSL / VAL.

**BCF** beat-cross frequency in Hz. The rate at which the curvilinear path crosses the average path. To calculate BCF we rotate each segment of h and A to 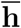 and 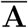 so that the rotated average path segment 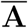 lies along the horizontal axis, sum the number of changes in sign in 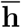 and divide by the time period.

**MAD** mean absolute angular displacement in degrees:

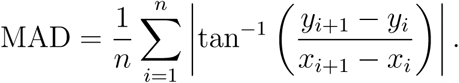

**TCS** we have also proposed the introduction of a new CASA measure for differentiating progressive cells, the track centroid speed, calculated as

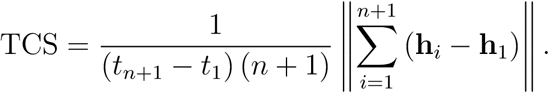

### Statistical analyses

Statistical analyses surrounding the relationship between directly measured flagellar beat frequency and beat cross frequency (BCF) consisted of line fitting in MATLAB, with the *R*^2^-values reported being the ratio of the sum of squares of the regression to the total sum of squares.

### Parameters for FAST

FAST parameters used in the analysis of the data in this manuscript can be found in the spreadsheets of data in the supplementary material.

### Ethical approval

All donors were recruited in accordance with the Human and Embryology Authority Code of Practice (Version 7) and gave informed consent (South Birmingham LREC 2003/239, and East Midlands REC 13/EM/0272).

## Results

The results of the analysis of experimental sets (1) and (2) (for some characteristic sperm) are shown for (1) free swimming cells in figures 1 and 2 (DSM and HVM respectively), and (2) adhered cells before and after addition of 4AP in figure 3.

**Figure 1:**
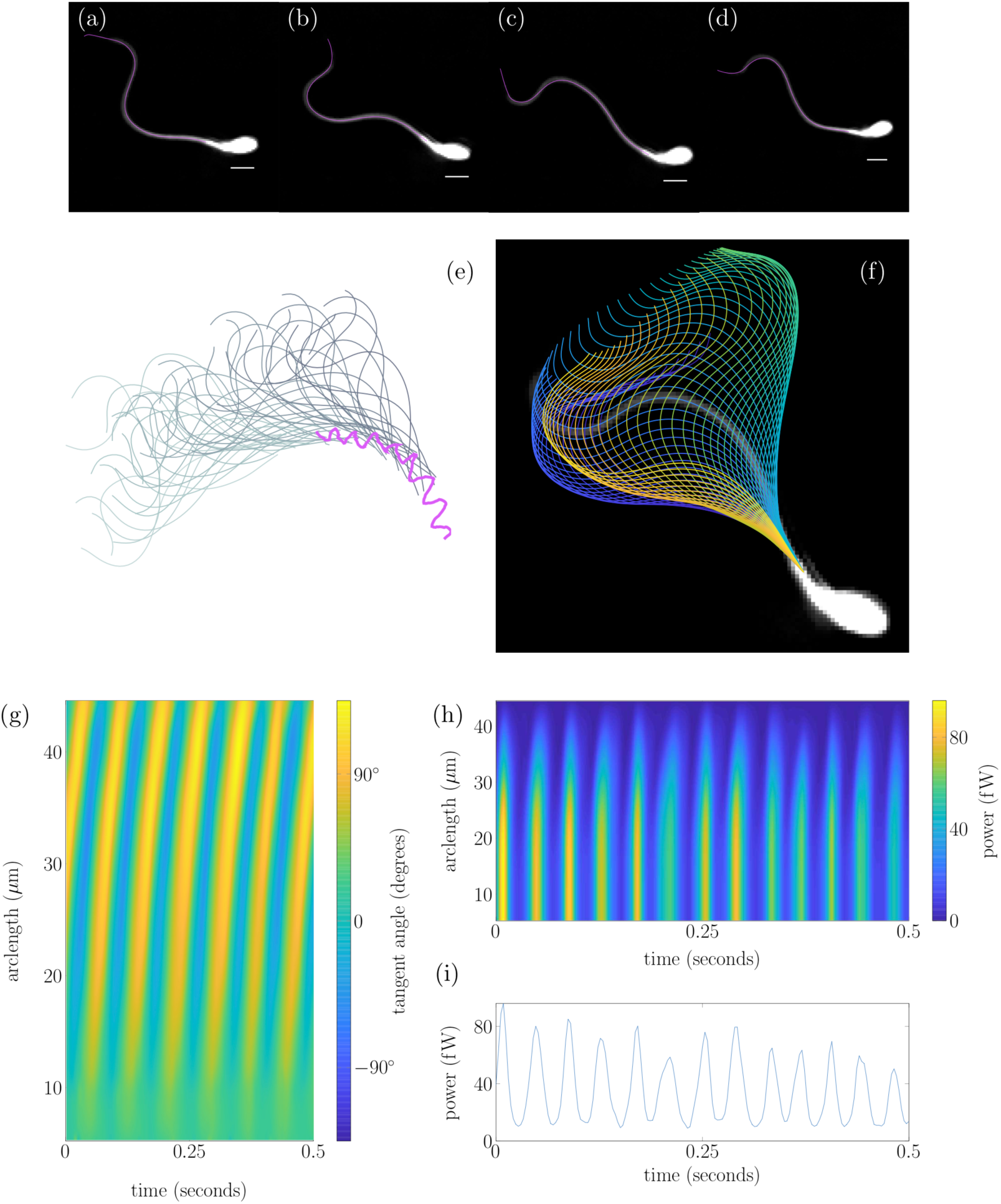
Tracking of a human sperm from experimental data set (1) in the diluted semen medium. Panels (a) – (d) show an overlay of the tracked flagellum over experimental frames at four points in a beat cycle, with a 5 *μ*m white scale bar. (e) shows the sperm head-track in magenta with associated flagellum plotted 0.014 s apart. (f) plots the flagellar beat over a single experimental frame with the colour of each flagellum representing time from dark blue to yellow. Panel (g) plots the tangent angle along the flagellum in the cell frame for 0.5 s. (h) shows the power exerted by the flagellum on the fluid distal to the point in arclength chosen, with the power exerted by the full flagellum shown in panel (i).

**Figure 2:**
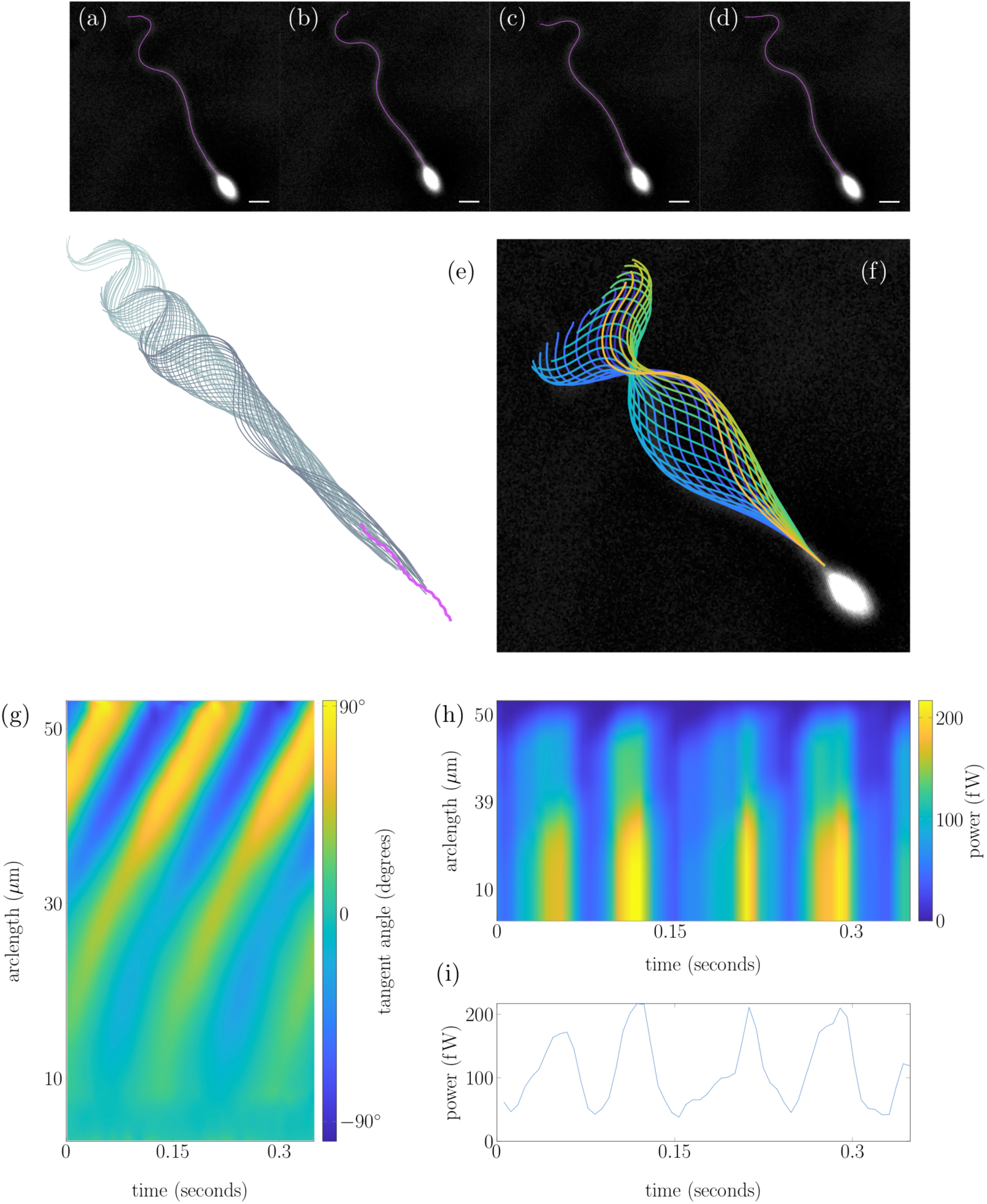
Tracking of a human sperm from experimental data set (1) in the high viscosity medium. Panels (a) – (d) show an overlay of the tracked flagellum over experimental frames at four points in a beat cycle, with a 5 *μ*m white scale bar. (e) shows the sperm head-track in magenta with associated flagellum plotted 0.007 s apart. (f) plots the flagellar beat over a single experimental frame with the colour of each flagellum representing time from dark blue to yellow. Panel (g) plots the tangent angle along the flagellum in the cell frame for 0.35 s. (h) shows the power exerted by the flagellum on the fluid distal to the point in arclength chosen, with the power exerted by the full flagellum shown in panel (i).

**Figure 3:**
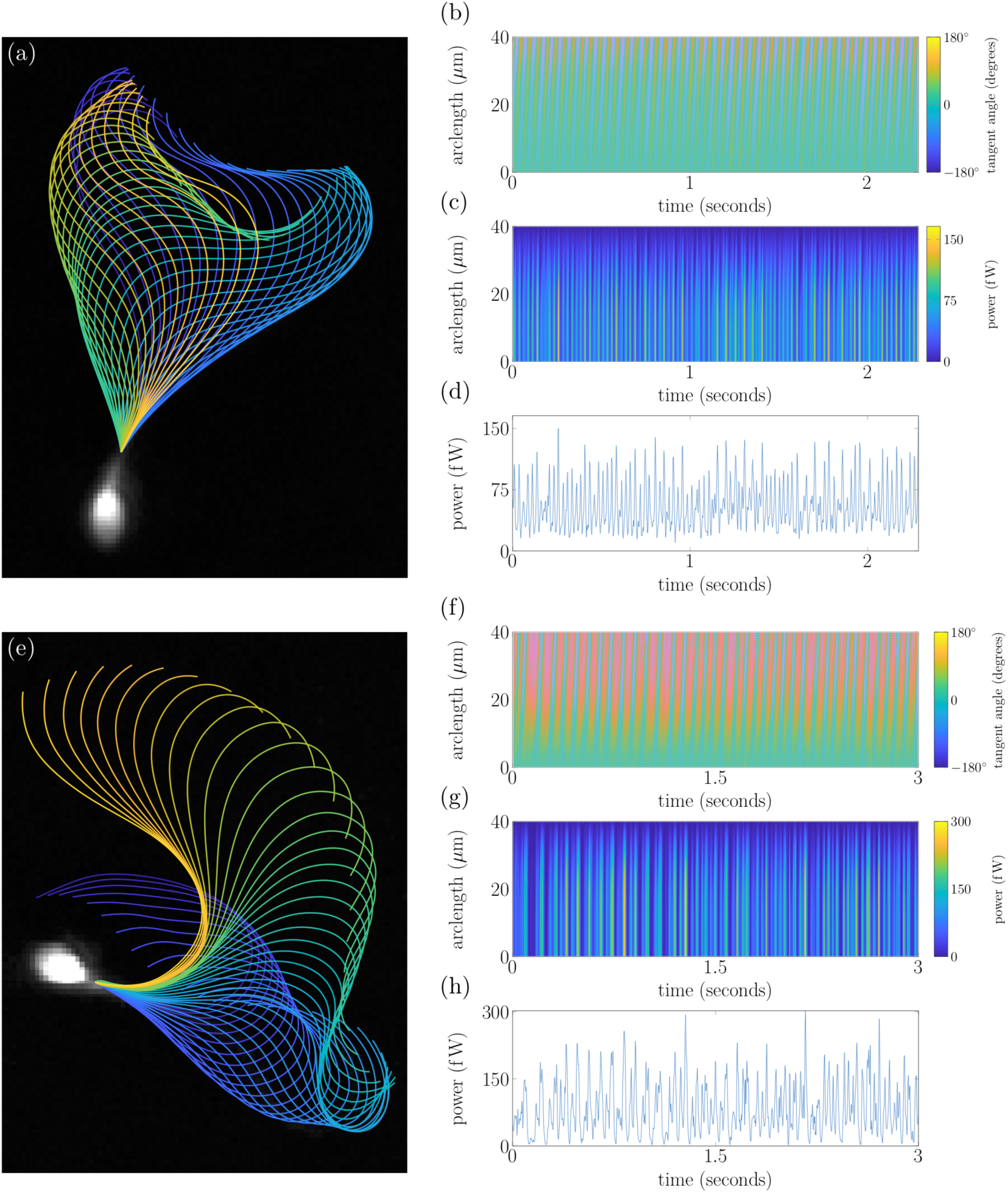
Tracking of stuck sperm from experimental data set (2) enables long-time analysis of cells. (a) – (d) the flagellar beat, tangent angle, power exerted by the flagellum distal to a point in arclength, and total power exerted by the flagellum respectively. (e) – (h) the same plots for a hyperactivated cell after stimulation with 4AP.

For experimental set (1) we plot λ, *f,* power dissipation, and three of the established CASA measures (curvilinear and average velocities (VCL, VAP) and beat cross frequency (BCF)) for each cell tracked in a scatter plot matrix (figure 4). The distinction between motilities in DSM and HVM are immediately apparent, forming two distinct sub-populations in the data. Notably, sperm in DSM have a median arc-wavelength of 31 *μ*m with IQR of 13 *μ*m, decreasing down to a median arc-wavelength of 17 *μ*m with IQR of 7 *μ*m in HVM. Similarly, cells in DSM have a median beat frequency of 19 Hz with IQR of 6.5 Hz, decreasing to a median beat frequency of 10 Hz with and IQR of 2.9 Hz in HVM. Note that in this manuscript we are using the Lighthill (1975) notion of arc-wavespeed and arc-wavelength rather than the conventional wavespeed and wavelength. This is due to the variation in the axis of wave propagation for sperm flagellum, making measurements in terms of the arclength along the flagellum a more natural choice.

**Figure 4:**
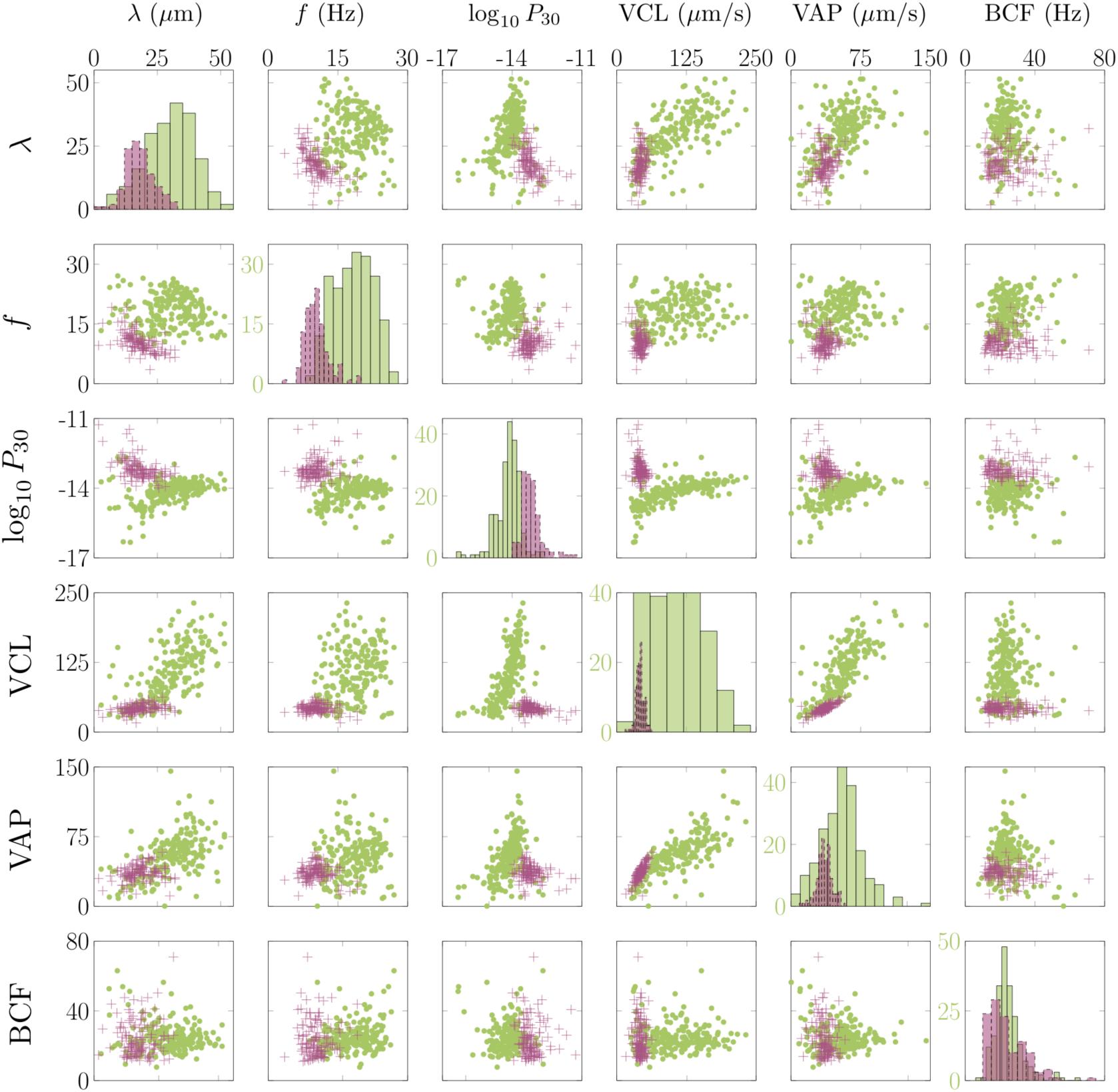
Scatter plot matrix showing relationships between arc-wavelength λ, flagellar beat frequency *f,* power generated by the first 30 *μ*m of flagellum *P*_30_ (measured in watts and plotted on a log-scale), the curvilinear velocity of the head VCL, the average path velocity of the head VAP and the beatcross frequency of the head BCF. Axes persist from left-to-right and top-to-bottom except on the leading diagonal where frequencies are shown in green. In each plot sperm swimming through HVM are shown as magenta crosses, and sperm swimming through DSM as green dots.

### Metabolic requirements of motility from imaging data

Knowledge of the flagellar waveform allows, via fluid dynamic modelling, for calculation of the power dissipation due to the beating of the flagellum. There are many methods to do this, each providing a balance between accuracy and ease-of-implementation (Gaffney et al., 2011; Gallagher and Smith, 2018; Smith, 2018). After analysis with FAST the captured flagellar movement can be used as input to a fluid mechanics calculation, which can then be used to assess the hydrodynamic power dissipation along the flagellum as demonstrated by Ooi et al. (2014). In the present work we treat the surrounding fluid as Newtonian, allowing the application of resistive force theory (Gray and Hancock, 1955; Gueron and Liron, 1992; Lighthill, 1976) (details in the methods section), which is acceptably accurate for the analysis of sperm motility (Friedrich et al., 2010; Johnson and Brokaw, 1979). Recently, modifications of resistive force theory have been suggested for non-Newtonian fluids (Riley and Lauga, 2017) which would enable the use of tracking data in modelling behaviours in more biologically complex fluids.

In figures 1i, 2i, 3d and 3h we plot the total power dissipation in watts integrated along the flagellum against time for a typical cell swimming in our low and high viscosity media, stuck, and stuck and stimulated with 4AP respectively. To understand how the power dissipation varies along the flagellum in figures 1h, 2h, 3c and 3g we plot the time-dependent power dissipation distal to a point in arclength along the flagellum. For fair comparison between cells when a varying length of flagellum is visible in experiments (often due to cell rolling or significant out-of-plane beating) we calculate the power dissipation by the first 30 *μ*m of flagellum (*P*_30_, shown with comparison to other measures in figure 4). The cells tracked in DSM reveal a median *P*_30_ of approximately 8.4 fW^1^ with interquartile range (IQR) of 9.4 fW. In contrast the cells swimming in HVM have an increased median *P*_30_ of 59 fW with IQR of 61 fW. For validation of the software package we have reanalysed the Ooi et al. (2014) data, who reported a median 7% decrease in cell power after the addition of 4AP; the reanalysis of a subset of these cells revealed a commensurate 19% decrease in the value of *P*_30_, where it should be noted exact comparisons of power dissipation over the full length of flagellum cannot be made due to the increased length of captured flagellum by FAST.

### Relationship between traditional CASA measures and sperm flagellar kinematics

In figure 4 we show comparisons between the flagellar arc-wavelength (λ), flagellar beat frequency (*f*), and the power dissipation through the first 30 *μ*m of flagellum (*P*_30_), as well as three of the commonly used existing CASA measures, namely the curvilinear velocity (VCL), average path velocity (VAP) and beat cross frequency (BCF). The change in media from DSM to HVM defines two clear clusters in the data.

#### BCF does not measure flagellar beat frequency

Beat cross frequency (BCF, the frequency with which the curvilinear path of the sperm head crosses the average path) was introduced as a proxy for flagellar beat frequency as it was impossible to make the correct measurement (Mortimer, 1997). We have been able to assess this quantitatively, with the results plotted in figure 4. Attempting to fit to the data we obtain *R*^2^-values of 0.042 and 0.00054 for cells in DSM and HVM respectively, showing that BCF does not measure the flagellar beat frequency.

#### Flagellar kinematics provide much stronger separation of different motility modes than CASA

Assessment of the data, as seen in the histograms in figure 4, reveals that the flagellar kinematic measures show greater separation between sperm in DSM vs HVM than standard CASA parameters. This not only emphasises the dramatic differences in sperm motility in different media, but also the importance of being able to accurately characterise them.

### New insights into sperm dynamics with modelling

The wealth of data provided by accurately tracking the sperm flagellar beat provides an exciting opportunity for use with mathematical models to gain new insights into the fluid dynamic properties induced by sperm swimming. Previously, to visualise the fluid velocity field surrounding a swimming cell required the use of experimental techniques such as micro-particle image velocimetry (Hamad, 2017), which is time consuming and prone to error. Instead we are able to pass an extracted flagellar beat, together with a model of a sperm head, to the NEAREST code package (Gallagher et al., 2018a; Gallagher and Smith, 2018; Smith, 2018). By solving the resulting fluid dynamics problem for a swimming sperm we are able to calculate the fluid velocity field surrounding the cell, plotted in figure 5. Here we see that there are significant differences in the velocity fields between the typical DSM and HVM cells, with the swimming of the cell in DSM inducing a much greater change in the fluid velocity due to the beating of the flagellum.

**Figure 5:**
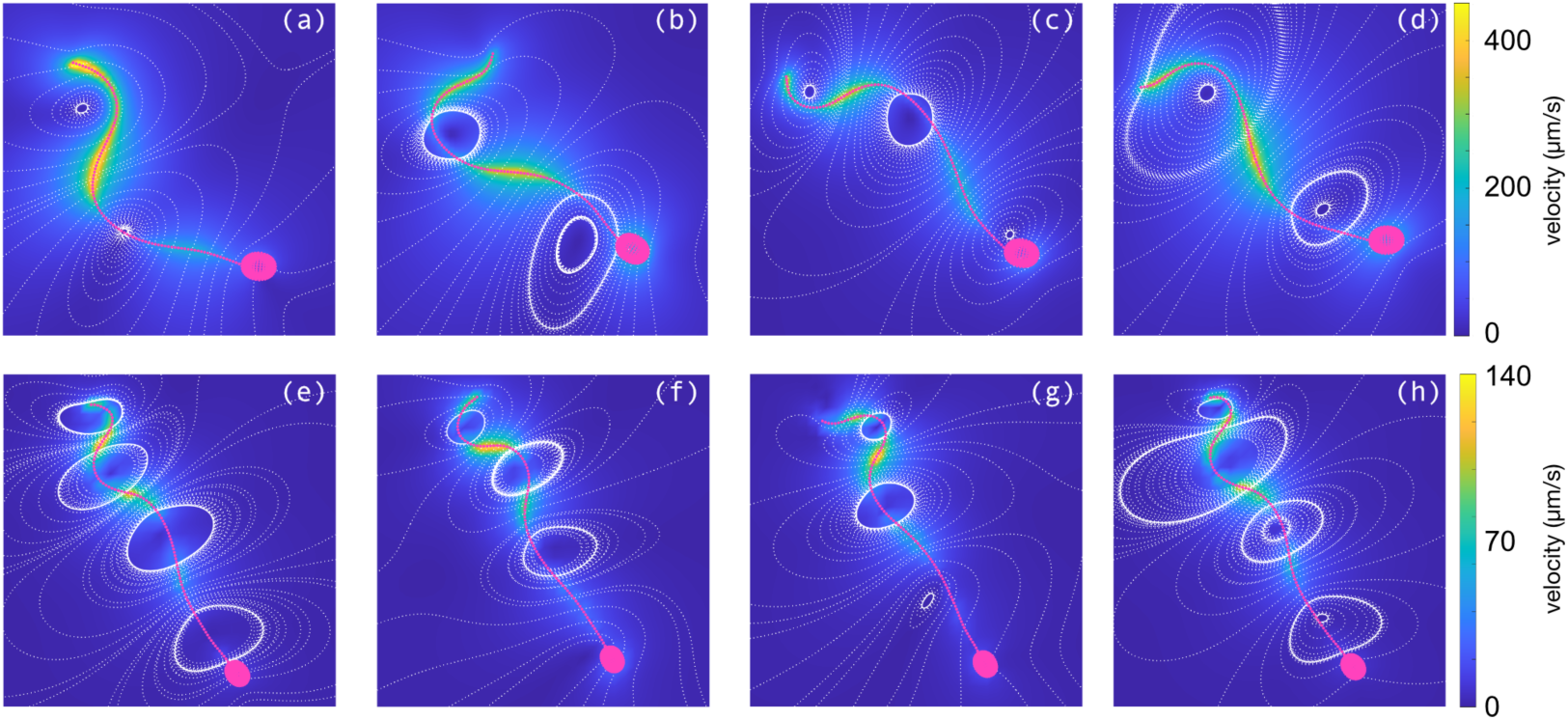
Simulated velocity fields with NEAREST for the tracked sperm in figures 1 and 2. The FAST tracked flagellum for each sperm has been paired with an idealised head and simulated in a 3D environment. Panels (a) – (d) show the flow fields in DSM at times corresponding to figures 1a – 1d, while (e) – (h) are in HVM, corresponding to 2a – 2d. In each figure the colour depicts the fluid velocity magnitude, with the sperm cell overlain in magenta and streamlines shown as white dots.

#### A new way to categorise progressive sperm

The WHO-IV (WHO, 1999) threshold for classifying a cell as progressive is a velocity greater than 5 *μ*m/s, however there is no consensus on which of the established CASA velocity measures should be used for this classification: VSL (straight-line velocity from first point tracked to last) may be misleading when sperm motility is circular as it has a significant dependence on the length of time the cell is tracked for, while the use of VCL and VAP may result in misclassification of cells which are twitching or being buffeted in tight circles as being progressive. To this end we propose a new measure, the track centroid speed (TCS, defined in the methods section), for this purpose. In figure 6 we plot the receiver operating characteristic (ROC) curves for characterising progressive sperm using TCS, as well as the standard CASA measures of VSL and VAP, against the gold standard of manual classification by a trained analyst. For each curve we see a significant area under the curve of between 0.964 and 0.971, with similar sensitivities of 98.1% to 99.7% (with TCS taking the lower of each of these measures). Where TCS truly outperforms the use of VSL and VAP is in the specificity of 85.6% compared to 78% and 60.6% for VSL and VAP respectively.

**Figure 6:**
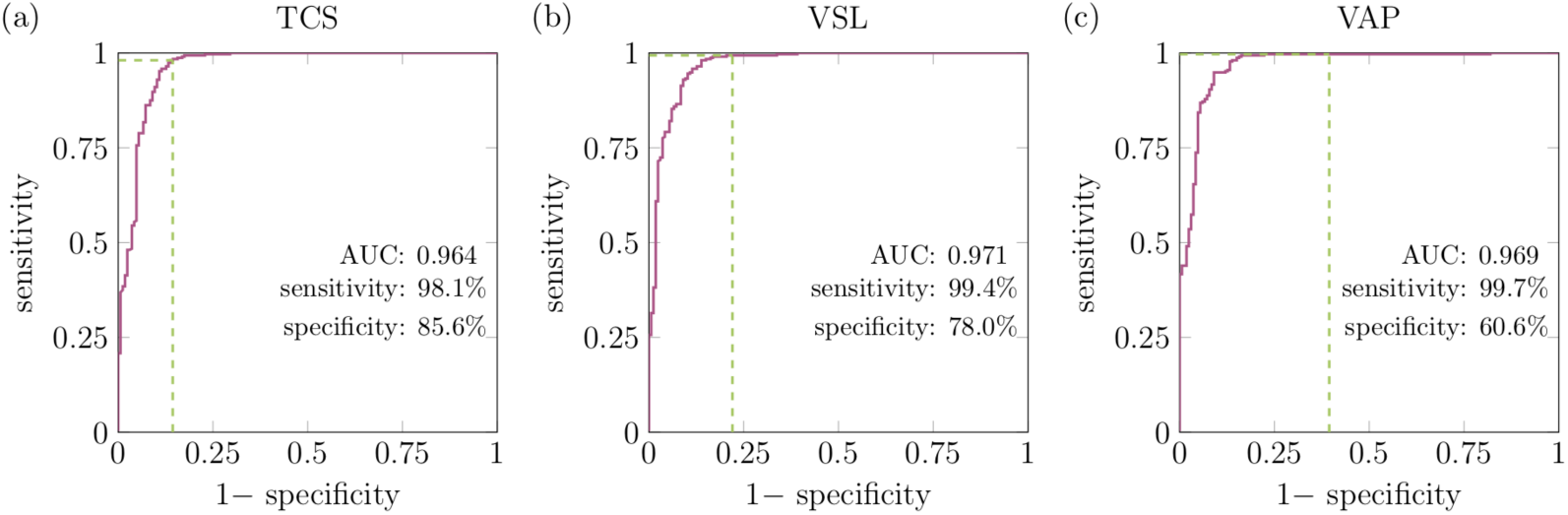
Receiver operating characteristic (ROC) curves for characterising sperm as progressive compared to the gold standard manual classification by a trained analyst. In panel (a) the ROC curve using TCS is plotted, with the more standard use of VSL and VAP shown for comparison in panels (b) and (c). The green lines highlight the sensitivity and specificity when using the 5 *μ*m/s WHO categorisation (WHO, 1999).

## Discussion

Meaningful clinical information in a heterogeneous population requires analysis of large cell numbers. Although the sperm flagellar waveform contains a wealth of potential diagnostic power, until now this could not be harnessed as the necessary tools were unavailable. This heterogeneity of human sperm kinematic motion means that excellent studies, which have been performed on small numbers of cells, have only been able to hint at underlying mechanisms and relationships, such as the link between sperm velocity and flagellar movement. Furthermore, there is a clear desire for new tools to assess large quantities of cells for human fertility research as well as male reproduction more generally. The FAST software package provides this ability, enabling assessment of large numbers of individual cells and the statistical power necessary to understand sperm motility on the population scale.

Rapid sperm motility is crucial for natural fertilisation (Hirano et al., 2001; Holt et al., 1985), and the diagnostic accuracy of this should improve with the availability of flagellar beat analysis. Developing a correct understanding of sperm motility will provide a potential diagnostic tool for male health more generally, and a substantial base for therapeutic developments. To address this we have developed and released FAST, the free-to-use software package for high-throughput flagellar beat analysis and cell tracking. We have tracked and analysed 366 sperm. By describing the flagellum as a tangent angle formulation we are able to provide detailed reporting of flagellar kinematic measures such as beat frequency and arc-wavelength for individual and populations of cells. In addition, the software provides improved accuracy for the existing CASA measures of motility. In particular, these data highlight that the existing BCF parameter is an incorrect extrapolation of flagellar beat frequency. In the present manuscript we have focussed on 2D imaging, exploiting the largely planar nature of the beat of the human sperm, however advances in imaging are increasingly making it possible to take 3D scans of swimming microorganisms (Pimentel et al., 2012; Su et al., 2012). It would be of interest, particularly for species other than human, to extend the capabilities of the software for non-planar flagellar motion.

Combining tracked flagella with mathematical modelling has the potential to reveal new mechanistic insight, for example it is now possible to estimate the metabolic requirements of motility a further quantity which is not accessible when only the head of a cell is tracked. Particle-based methods to image flow on microscopic scales are technically challenging and limited in resolution due to Brownian effects. In contrast, the combination of negative phase contrast microscopy, FAST and the NEAREST fluid dynamics package (Gallagher and Smith, 2018) provide a much more convenient, rapid and highly-resolved (if indirect) method to estimate fluid dynamic effects, such as that of a swimming cell on its surrounding environment. We have designed FAST to be agnostic to fluid mechanics, enabling the coupling of tracked analysis with more advanced computational methods – for example incorporating non-Newtonian effects – as they become available.

The novel ability to quantify accurate flagellar beat detail in large populations of motile cells enables an abundant array of diagnostic, toxicological, and therapeutic possibilities. In particular we hope that it opens the way for new approaches to assessing and treating male subfertility.

## Data accessibility

The FAST software package and all documentation can be downloaded from www.flagellarCapture.com.

## Author’s roles

M.T.G. designed and implemented the algorithms, wrote the paper, collected and analysed data, designed the research. G.C. collected and analysed data, co-wrote the paper, contributed to algorithm design and testing. E.H.O. performed adhered cell imaging experiments. J.C.K-B. conceived the research, co-supervised the project, co-wrote the paper, led the clinical concepts and interpretation, and supervised laboratory work. D.J.S. conceived the research, supervised the project, co-wrote the paper, contributed to algorithm design, and co-supervised laboratory work.

## Acknowledgements

The ongoing support of patients and staff at the Birmingham Women’s and Children’s NHS Trust are fundamental to our research work.

## Footnotes

1 One femtowatt (fW) is equal to 10^-15^ watts.

## References

Amann, R., and Waberski, D. Computer-assisted sperm analysis (CASA): capabilities and potential developments. Theriogenology 2014; 81, 5–17.

Friedrich, B., Riedel-Kruse, I., Howard, J., and Jülicher, F. High-precision tracking of sperm swimming fine structure provides strong test of resistive force theory. J. Exp. Biol. 2010; 213, 1226–1234.

Gaffney, E., Gadêlha, H., Smith, D., Blake, J., and Kirkman-Brown, J. Mammalian sperm motility: observation and theory. Annu. Rev. Fluid Mech. 2011; 43, 501–528.

Gallagher, M., Choudhuri, D., and Smith, D. Sharp quadrature error bounds for the nearest-neighbor discretization of the regularized stokeslet boundary integral equation. arXiv preprint arXiv:1806.01560 2018a;.

Gallagher, M., and Smith, D. Meshfree and effcient modeling of swimming cells. Phys. Rev. Fluids 2018; 3, 053101.

Gallagher, M., Smith, D., and Kirkman-Brown, J. CASA: tracking the past and plotting the future. Reprod. Fert. Develop. 2018b; 30, 867–874.

Gray, J., and Hancock, G. The propulsion of sea-urchin spermatozoa. J. Exp. Biol. 1955; 32, 802–814.

Gueron, S., and Liron, N. Ciliary motion modeling, and dynamic multicilia interactions. Biophys. J. 1992; 63, 1045.

Guerrero, A., Nishigaki, T., Carneiro, J., Tatsu, Y., Wood, C., and Darszon, A. Tuning sperm chemotaxis by calcium burst timing. Dev. Biol. 2010; 344, 52–65.

Hamad, A. Molecular and physical interactions of human sperm with female tract secretions. Ph.D. thesis University of Birmingham 2017.

Hansen, J., Rassmann, S., Jikeli, J., and Wachten, D. SpermQ-a simple analysis software to comprehensively study flagellar beating and sperm steering. bioRxiv 2018; (p. 449173).

Hiramoto, Y., and Baba, S. A quantitative analysis of flagellar movement in echinoderm spermatozoa. J. Exp. Biol. 1978; 76, 85–104.

Hirano, Y., Shibahara, H., Obara, H., Suzuki, T., Takamizawa, S., Yam-aguchi, C., Tsunoda, H., and Sato, I. Andrology: Relationships between sperm motility characteristics assessed by the computer-aided sperm analysis (casa) and fertilization rates in vitro. J. Assist. Reprod. Gen. 2001; 18, 215–220.

Holt, W., Cummins, J., and Soler, C. Computer-assisted sperm analysis and reproductive science; a gift for understanding gamete biology from multidisciplinary perspectives. Reprod. Fert. Develop. 2018; 30, iii–v.

Holt, W., Moore, H., and Hillier, S. Computer-assisted measurement of sperm swimming speed in human semen: correlation of results with in vitro fertilization assays. Fertil. Steril. 1985; 44, 112–119.

Van der Horst, G. IVF including ICSI needs CASA sperm functionality more than ever before! Microptic S.L. Blog 2017; URL: https://www.micropticsl.net/wordpress/ivf-including-icsi-needs-casa-sperm-functionality-more-than-ever-before-2/, accessed August 2017.

Inaba, K., and Shiba, K. Microscopic analysis of sperm movement: links to mechanisms and protein components. Microscopy; 2018; 67, 144–155.

Inhorn, M., and Patrizio, P. Infertility around the globe: new thinking on gender, reproductive technologies and global movements in the 21st century. Hum. Reprod. Update 2015; 21, 411–426.

Johnson, R., and Brokaw, C. Flagellar hydrodynamics. a comparison between resistive-force theory and slender-body theory. Biophys. J. 1979; 25, 113–127.

Kaneko, T., Mori, T., and Ishijima, S. Digital image analysis of the flagellar beat of activated and hyperactivated Suncus spermatozoa. Mol. Reprod. Dev. 2007; 74, 478–485.

Lighthill, M. Mathematical biofluiddynamics. SIAM 1975.

Lighthill, M. Flagellar hydrodynamics. SIAM review 1976; 18, 161–230.

Mortimer, S. A critical review of the physiological importance and analysis of sperm movement in mammals. Hum. Reprod. Update 1997; 3, 403–439.

Mukhopadhyay, A., and Dey, C. Reactivation of flagellar motility in demem-branated Leishmania reveals role of cAMP in flagellar wave reversal to ciliary waveform. Sci. Rep. 2016; 6, 37308.

Ohmuro, J., and Ishijima, S. Hyperactivation is the mode conversion from constant-curvature beating to constant-frequency beating under a constant rate of microtubule sliding. Molecular Reproduction and Development: Incorporating Gamete Research 2006; 73, 1412–1421.

Ooi, E., Smith, D., Gadêlha, H., Gaffney, E., and Kirkman-Brown, J. The mechanics of hyperactivation in adhered human sperm. Roy. Soc. Open Sci. 2014; 1, 140230.

Pimentel, J., Carneiro, J., Darszon, A., and Corkidi, G. A segmentation algorithm for automated tracking of fast swimming unlabelled cells in three dimensions. J. Microsc. 2012; 245, 72–81.

Reddy, G., Mukhopadhyay, A., and Dey, C. Characterization of ciliobrevin A mediated dynein ATPase inhibition on flagellar motility of Leishmania donovani. Mol. Biochem. Parasitol. 2017; 214, 75–81.

Riley, E., and Lauga, E. Empirical resistive-force theory for slender biological filaments in shear-thinning fluids. Phys. Rev. E 2017; 95, 062416.

Shiba, K., Tagata, T., Ohmuro, J., Mogami, Y., Matsumoto, M., Hoshi, M., and Baba, S. Peptide-induced hyperactivation-like vigorous flagellar movement in starfish sperm. Zygote 2006; 14, 23–32.

Smith, D. A nearest-neighbour discretisation of the regularized stokeslet boundary integral equation. J. Comp. Phys. 2018; 358, 88–102.

Smith, D., Gaffney, E., Gadêlha, H., Kapur, N., and Kirkman-Brown, J. Bend propagation in the flagella of migrating human sperm, and its modulation by viscosity. Cytoskeleton 2009; 66, 220–236.

Smith, D., Montenegro-Johnson, T., and Lopes, S. Symmetry-breaking cilia-driven flow in embryogenesis. Annu. Rev. Fluid Mech. 2019; 51, 105–128.

Su, T.-W., Xue, L., and Ozcan, A. High-throughput lensfree 3d tracking of human sperms reveals rare statistics of helical trajectories. P. Natl. Acad. Sci. 2012; 109, 16018–16022.

Walker, B., and Wheeler, R. Identifying flagella from videomicroscopy: exploiting a conserved morphology. arXiv preprint arXiv:1808.01249 2018.

Wan, K., and Goldstein, R. Time irreversibility and criticality in the motility of a flagellate microorganism. Phys. Rev. Lett. 2018; 121, 058103.

WHO WHO laboratory manual for the examination of human semen and sperm-cervical mucus interaction. (4th ed.). Cambridge university press 1999.

WHO WHO Laboratory Manual for the Examination and Processing of Human Semen, 5th ed. World Health Organisation 2010.

Wilson, L., Carter, L., and Reece, S. High-speed holographic microscopy of malaria parasites reveals ambidextrous flagellar waveforms. P. Natl. A. Sci. 2013; 110, 18769–18774.

Xu, H., Medina-Sánchez, M., Magdanz, V., Schwarz, L., Hebenstreit, F., and Schmidt, O. Sperm-hybrid micromotor for targeted drug delivery. ACS nano 2017; 12, 327–337.

WHO WHO laboratory manual for the examination of human semen and sperm-cervical mucus interaction. (4th ed.).Cambridge university press 1999.

